# Deep learning of antibody epitopes using molecular permutation vectors

**DOI:** 10.1101/2024.03.20.585661

**Authors:** Ioannis Vardaxis, Boris Simovski, Irantzu Anzar, Richard Stratford, Trevor Clancy

## Abstract

**Background:** The accurate computational prediction of B cell epitopes can vastly reduce the cost and time required for identifying potential epitope candidates for the design of vaccines and immunodiagnostics. However, current computational tools for B cell epitope prediction perform poorly and are not fit-for-purpose, and there remains enormous room for improvement and the need for superior prediction strategies.

**Results:** Here we propose a novel approach that improves B cell epitope prediction by encoding epitopes as binary molecular permutation vectors that represent the position and structural properties of the amino acids within a protein antigen sequence that interact with an antibody, rather than the traditional approach of defining epitopes as scores per amino acid on a protein sequence that pertain to their probability of partaking in a B cell epitope antibody interaction. In addition to defining epitopes as binary molecular permutation vectors, the approach also uses the 3D macrostructure features of the unbound 3D protein structures, and in turn uses these features to train another deep learning model on the corresponding antibody-bound protein 3D structures. We demonstrate that the strategy predicts B cell epitopes with improved accuracy compared to the existing tools. Additionally, we demonstrate that this approach reliably identifies the majority of experimentally verified epitopes on the spike protein of SARS-CoV-2 not seen by the model in training and generalizes in very robust manner on dissimilar data not seen by the model in training.

**Conclusions:** With the approach described herein, a primary protein sequence with the query molecular permutation vector alone is required to predict B cell epitopes in a reliable manner, potentially advancing the use of computational prediction of B cell epitopes in biomedical research applications.

## INTRODUCTION

B-cell epitopes (BCEs) are clusters of surface accessible amino acids on a protein antigen, recognized by B cell secreted antibodies or B cell receptors (BCR) (1). The B cell molecular recognition of BCEs by BCRs elicits humoral and cellular immune responses that are key in the fight against pathogenic threats. Knowledge of the precise coordinates of antibody epitope contact points in an antigen upon binding to an antibody can be of tremendous value. Such BCE information can offer crucial guidance in vaccine design (2, 3), therapeutic antibody engineering (2), in the streamlining of numerous diagnostic (4, 5), and therapeutic applications in molecular medicine (4). Hence, a variety of BCE mapping strategies have been developed to identify such clusters of BCE coordinates on antigens. Many of these are wet lab-based methods such as X-ray co-crystallography, cryogenic electron microscopy (cryo-EM) and numerous other assays (6). However, there are innumerable possible BCEs harbored on any given protein antigen sequence, and the experimental approaches to capture these are extremely time consuming, laborious, and expensive, and therefore not amenable to be applied on a large-scale for comprehensive BCE mapping and screening. The ability to accurately predict BCEs computationally would greatly facilitate the comprehensive mapping of complex antigens helping to speed up the development of vaccine and immune-based diagnostics (6, 7). In recent years, numerous computational prediction algorithms have been developed to attempt the *in-silico* BCE mapping of protein antigens (7, 8). However, the accurate computational prediction of BCEs remains a daunting challenge as most of these prediction tools perform poorly in comparative benchmarking studies (7, 9, 10) and are consequently not fit for purpose. However, successfully addressing the current limitations in the computational prediction of BCEs has the potential to revolutionize vaccine and immunodiagnostics development (11). Here we propose a novel approach to BCE prediction that significantly improves the predictive performance compared to the current state-of-the-art tools. Our approach encompasses several novel and distinguishing innovations that more accurately model the underlying biology of BCE/antibody recognition. These include, (1) the development of a set of Bidirectional Long Short-Term Memory (BLSTM) models that predict pertinent 3D features of the protein antigen (we term here as «3D macrostructure» features), thereby circumventing the need for experimentally derived or computationally predicted 3D protein structures, (2) predicting the aforementioned 3D macrostructure features based on the “unbound” rather than the antibody “bound” protein antigen structure, and (3) defining or encoding the epitopes used for training as binary molecular permutation vectors (MPVs) that represent the 3D physical interaction of the BCE with an antibody. We demonstrate here that the proposed strategy outperforms existing conformational BCE prediction approaches and successfully predicts the majority of unseen SARS-CoV-2 experimentally verified epitopes.

## RESULTS

### Outline for a high performing B cell epitope predictor

A top-level overview of the BCE-HUNT approach to epitope prediction is illustrated in *Figure 1*. The high performance we report in the subsequent sections below, delivered by BCE-HUNT, is achieved by the two main innovations that distinguish this approach compared to the state-of-the-art BCE predictors. Firstly, the approach circumvents the need for experimentally derived 3D protein structures and/or computationally predicted 3D protein structures (which are not yet fit for purpose(12)), by using protein sequence features and numerous other physiochemical features to predict 3D macrostructure properties (see Table 1 for the complete list of features and their sources). The 3D macrostructure properties are defined here as the key determinants of protein surface exposure of the protein sequence regions, namely, relative solvent accessibility (RSA), upper half-sphere exposure (HSE), lower half-sphere exposure (LHSE), and secondary structure (SS), as outline in *Figure 1A*. Critically, in this first innovation, the 3D macrostructure features are learned from the unbound 3D protein structures (Figure 1A). The pipeline trains distinct BLSTM deep learning models for each of the 3D macrostructure properties to predict these properties from all the relevant unbound single 3D protein structure features in the Protein Data Bank (PDB) (13). The output at prediction time for each of these BLSTM models is a score for each amino acid of the input protein sequence for each of the 3D macrostructure properties, thereby negating the need for knowing or predicting the exact coordinates of each atom in each amino acid relative to each other offered by complete and accurate experimental 3D protein structure determination (see *Figure 1A*). Secondly, the proposed strategy relies on the redefinition of an epitope in BCE predictors by not only scoring an epitope based on a per amino acid basis in the protein sequence, but rather by molecular permutation vector (MPV) of amino acids. In MPVs 1’s represents direct amino acid contact points on the 3D protein sequence with the antibody, and 0’s represents amino acids on the protein sequence that are not interacting directly with the antibody *(see Figure 1B)*. This differs from the existing state-of-the-art predictors which primarily score each amino acid (AA) on a per AA basis for its potential contribution to the BCE interaction with the antibody. In the MPV definition of an epitope the entire BCE interaction structure is defined and predicted. Critically, in this second innovation, the BCE interaction structure is learned from the antibody-bound 3D protein complex structures in the PDB (Figure 1B). These two innovations form the basis of the main BCE-HUNT predictor. As outlined in *Figure 1B,* similar to the 3D macrostructure predictors, the main BCE predictor model (BCE-HUNT) is also trained using BLSTMs, but in this step the training data is based on all the antibody-bound protein structure complexes from the PDB using the prediction output of the 3D macrostructure features in *Figure 1A* in addition to the proteins sequence features (additional features are also used, see Table 1). At prediction time the input to the main BCE-HUNT predictor model requires only the sequence, and the query MPV (sequence vector of 1’s and 0’s represent the protein sequence, where 1 represents an amino acid contact point with the antibody and 0 represent not contact with the antibody). The main output for BCE-HUNT is a score ranging from 0 to 1.0, representing the probability that the MPV for the primary protein sequence being queried is a true positive BCE. The BCE-HUNT complete pipeline workflow for the prediction of both linear and conformational BCE by BCE-HUNT is outlined in Supplementary Figure 1.

**Figure 1:**
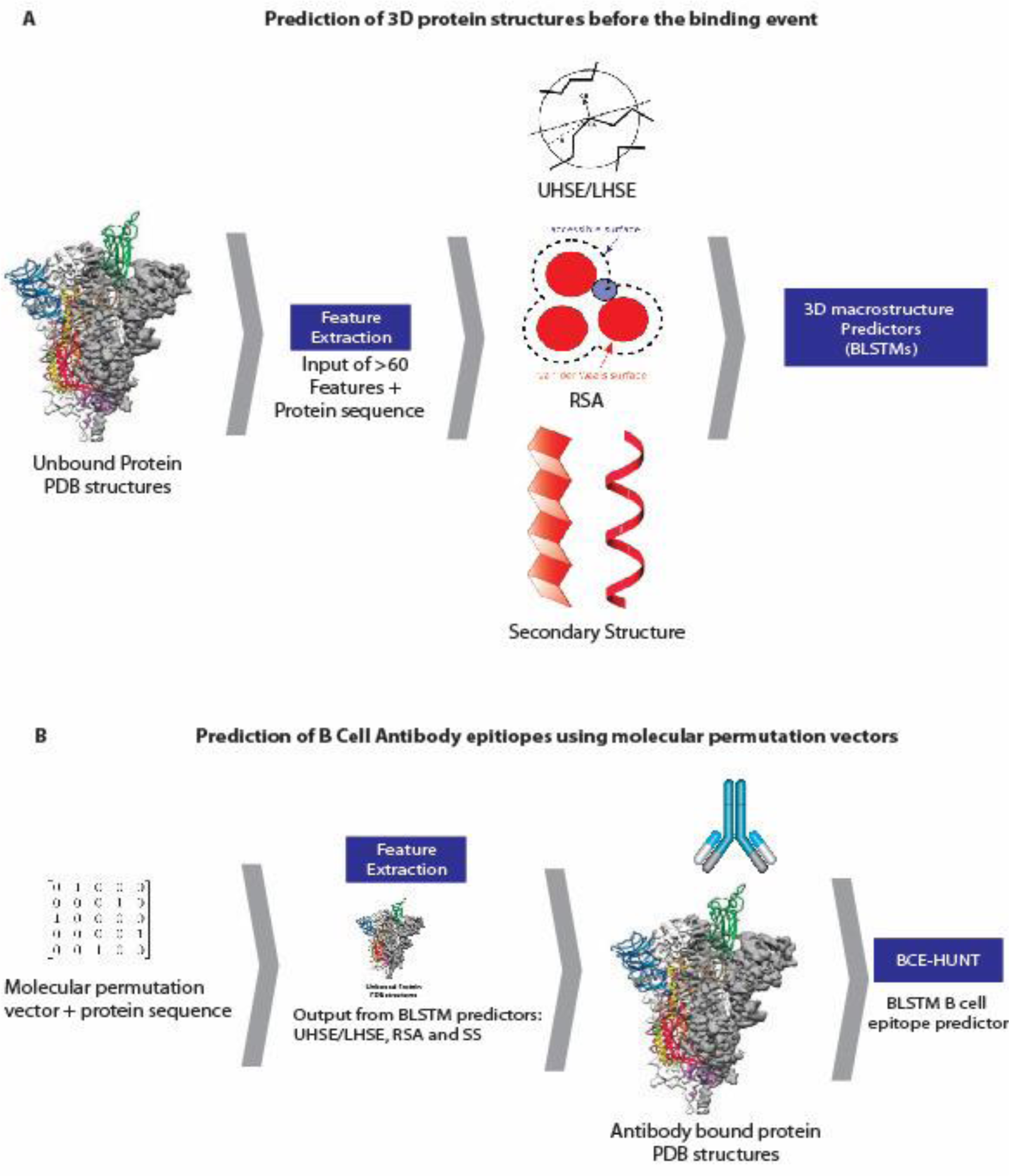
BCE Hunt overview. A top-level outline of the two-step innovative pipeline underlying BCE-HUNT. In step 1, depicted in Figure 1A a BLSTM based deep learning predictor is trained based on all single 3D protein structures that are unbound (*i.e.,* 3D protein structures not bound to any antibody or other ligand) in the PDB database. This predictor learns the 3D macrostructure determinants of 3D protein folding. Namely, RSA, UHSE and LHSE. The features used to train these 3D macrostructure models are derived from the unbound 3D protein structure sequences in addition to numerous other amino acid physiochemical and 3D protein structure sequences (see Table 1). In step 2, depicted in Figure 1B a BLSTM based deep learning predictor is trained based on all of the relevant antibody-bound 3D protein structure complexes in the PDB. The features used in training are in essence the protein sequences of the antibody-bound protein structure and the predicted 3D macrostructure protein features for the unbound protein structure, predicted as described in Figure 1A. The input at prediction for the main BCE model here is the target antigen protein sequence, and the query molecular permutation vector (MPV) that defines the antibody BCE (sequence vector of 1’s and 0’s represents the protein sequence, where 1 represents an amino acid contact point with the antibody and 0 represent not contact with the antibody). Full details are described in methods and the complete ML training pipeline outlined in Supplementary Figure 1.

**Table 1:** Features used to train the 3D macrostructure predictors – Upper half sphere exposure (UHSE), Lower half sphere exposure (LHSE), Relative Solvent Accessibility (RSA), and secondary structure (SS). In addition to the features used to train the final main BCE prediction model, BCE-HUNT. The source information for each feature is referenced where relevant.

| UHSE, LHSE & RSA | SS | BCE-HUNT |
| --- | --- | --- |
| Polarity (14) | Conformational parameter for coil (15) | Bulkiness (16) |
| Free energy of transfer from inside to outside of a globular protein (17) | Average surrounding hydrophobicity (18) | Polarity (14) |
| Hydrophobicity (19) | Conformational preference for total beta strand (antiparallel and parallel) (20) | Molar fraction of buried residues (17) |
| Membrane buried helix parameter (21) | Normalized frequency for beta-sheet (22) | Conformational parameter for beta-turn (23) |
| Hydrophobicity (24) | Conformational parameter for alpha helix (15) | Average flexibility index (25) |
| Transmembrane tendency (26) | Side chain classes (27) | Normalized frequency for beta-sheet (22) |
| Proportion of residues buried (28) | Normalized frequency for beta-turn (22) | Normalized frequency for alpha helix (22) |
| Mean fractional area loss (29) | Binary representation of protein sequence | Side chain classes (27) |
| Conformational parameter for beta-sheet (23) |  | Known Number of codon(s) coding for each amino-acid in universal genetic code (27) |
| Conformational preference for total beta strand (20) |  | Side chain polarity (27) |
| Side chain classes (27) |  | Binary representation of protein sequence |
| Surface accessibility (30) |  | Secondary Structure (internal algorithm) |
| Side chain polarity (27) |  | Relative Solvent Accessibility (internal algorithm) |
| Binary representation of protein sequence |  | Molecular Permutation Vectors |

### Evaluation metrics of the main BCE-HUNT BLSTM model: cross validation and independent tests

For both the 3D macrostructure BLSTM models and the main BCE BLSTM model (BCE-HUNT) we used∼ 80% of the training data on 5-fold cross-validation (CV) to assess the performance of the models, and the remaining 20% was used as independent test against the trained models. All the models performed well, without overfitting, and had high performance for all evaluation metrics. Moreover, all the models demonstrated high stability, with only a small variation observed between different CV runs. In particular, the model performance for the main BCE predictor is illustrated in Figure 2. The model performed with a high precision-recall (PR) AUC of 0.8 compared the no skill value of 0.03 (robust across all CVs in Figure 2A-2E and comparable to the independent test in Figure 2F. This BLSTM model predicts conformational BCEs trained on the antibody-bound protein 3D structures, with input features from the 3D macrostructure models of the corresponding unbound protein 3D structures (depicted in Figure 1A and Supplementary Figure 1B), with the definition of the epitope based on MPVs. Although all the BLSTM models had promising and robust CV runs, and model training and independent tests exhibited high and robust performance, we next proceeded to evaluate the models against existing state-of-the-art BCE predictors.

**Figure 2:**
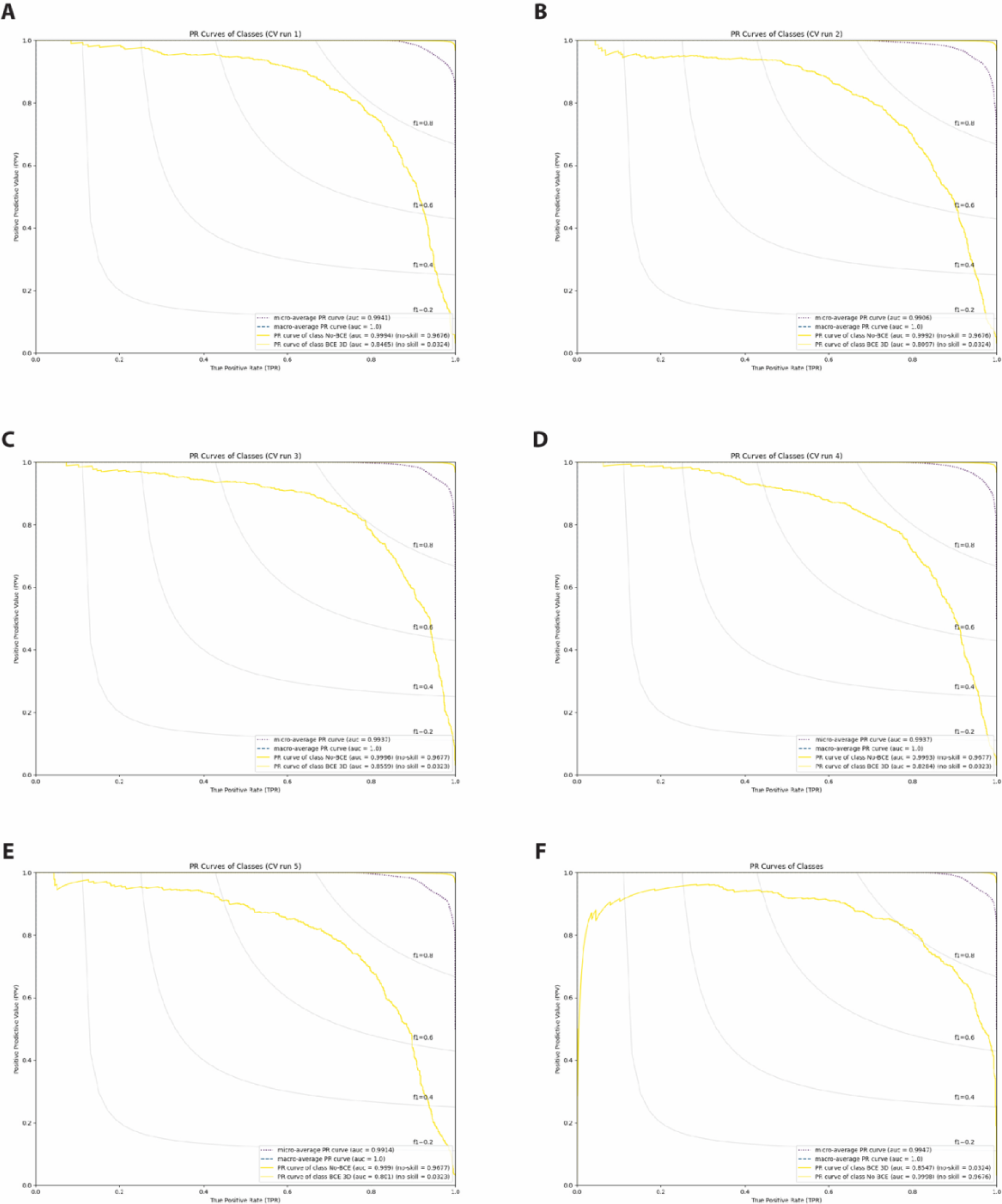
Cross validation and independent tests for the BCE-HUNT BLSTM model. Panels A-E on this figure shows results for the precision-recall (PR) metric for cross validation runs 1-5, respectively. PR was chosen for this evaluation due to the unbalanced nature of the dataset in terms of positives and negatives in the training data (see methods). The BCE-HUNT model exhibited a very high performance when using the no-skill PR metric. The average PR AUC for each of the CV runs 1-5 (panel A-E) was 0.83 compared to an average no skill PR of 0.03. The CV models (A-E) were trained for 459 epochs and independent test (F) for 583. In panel F we demonstrate the performance of the BCE-HUNT model on an independent, left out and non-redundant, test data set of PDB antibody protein binding structures. The performance in panel F illustrates that the BCE-HUNT model performs well on data not seen during training of the BLSTM model, the PR AUC curve depicts a robust and comparable performance compared to the CV results shown in A-E. “No-BCE” on the graphs are models trained on random positive and negative data.

Although the BLSTM and pipeline architecture of BCE-HUNT is setup for both linear and conformational BCEs, the current version is trained on conformational BCEs only. Based on the relevance for conformational BCE predictions, and operational availability for comparisons, three existing state-of-the-art tools were used to compare against, namely; Graphbepi (31), Discotope3.0 (32) and CBtope (33). Each of these three different tools have different scoring and predictions systems and the scores per AA was interpreted individually for each tool. The data set used in these comparisons was an independent test dataset, not used in the training for BCE-HUNT. For this independent test we used 6% of the most recent antibody-bound protein 3D structures in the PDB, and 94% of the PDB data was used on training (to avoid test overlaps with the other algorithms).

Benchmarking against existing state-of-the-art conformational BCE tools is challenging due to (1) their unique epitope definitions, based on different Armstrong (Å) distances between the epitope and the paratope contact points on the antibody in the 3D structures of antibody-protein antigen complex, and (2) their per amino acid (AA) BCE contribution prediction, in contrast to our method. That is, in the existing conformational BCE prediction tools each AA in the query protein sequence is scored according to its potential to contribute to a BCE antibody interaction. However, our tool BCE-HUNT conceptually defines an epitope based on MPVs representing the protein sequence, whereby we predict the entire epitope direct contact points represented as a permutation vector for each individual query protein sequence. In BCE-HUNT a probability for each protein sequence and single MPV is then outputted per query at prediction time. Therefore, for BCE-HUNT, the metrics are defined as predicting the entire epitope’s direct contact points and non-contact points with the antibody, whereby the existing tools assign a score for each AA on a protein sequence representing its potential contribution to participating in the BCE. This conceptual difference makes it challenging to directly compare BCE-HUNT against existing tools. However, to adjust for these conceptual differences, we made alterations to the architecture of BCE-HUNT such that in addition to outputting the probability scores of the MPV, the model also outputs a probability score per AA (to make comparisons against the existing tools possible).

In a first evaluation against the existing tools an experimental comparison was devised that is conceptually similar as to how an epitope is defined as MPVs in BCE-HUNT (Figure 3A). Here, the entire epitope needs to be predicted positive for it to count in any of the tools being analyzed (meaning each AA contributing experimentally validated to directly partake in the epitope-paratope interaction is predicted correctly). For example, if a true epitope in the test is represented by the following permutation vector [**1**,0,0,**1**,0,**1**] (where the 1’s represent experimentally verified direct contact points with the antibody at a predefined Å distance, and 0’s represent no direct contact with the antibody in the experimentally verified 3D structure). Therefore, for example, as Discotope3.0 reports in its model description that an AA contributes to the epitope if it has a score > 0.9 (32); the following was counted as a successful TP prediction for Discotope3.0 when considering the same permutation vector: [**0.95**,0.02,0.3,**0.98**,0.2,**0.91**]. For this evaluation we allowed the existing tools to count a successful TP when some of the 0’s in the epitope defined by the permutating vector above were also counted as positives. So, for example, based on the same > 0.9 threshold for Discotope3.0, the permutation vector [**0.95**,0.99,0.91,**0.98**,0.93,**0.91**] would also be counted as a successful TP prediction (but not for BCE-HUNT, more strictly, it was counted as a failure if one of the 0’s is predicted as an epitope contact point in Figure 3A).

**Figure 3:**
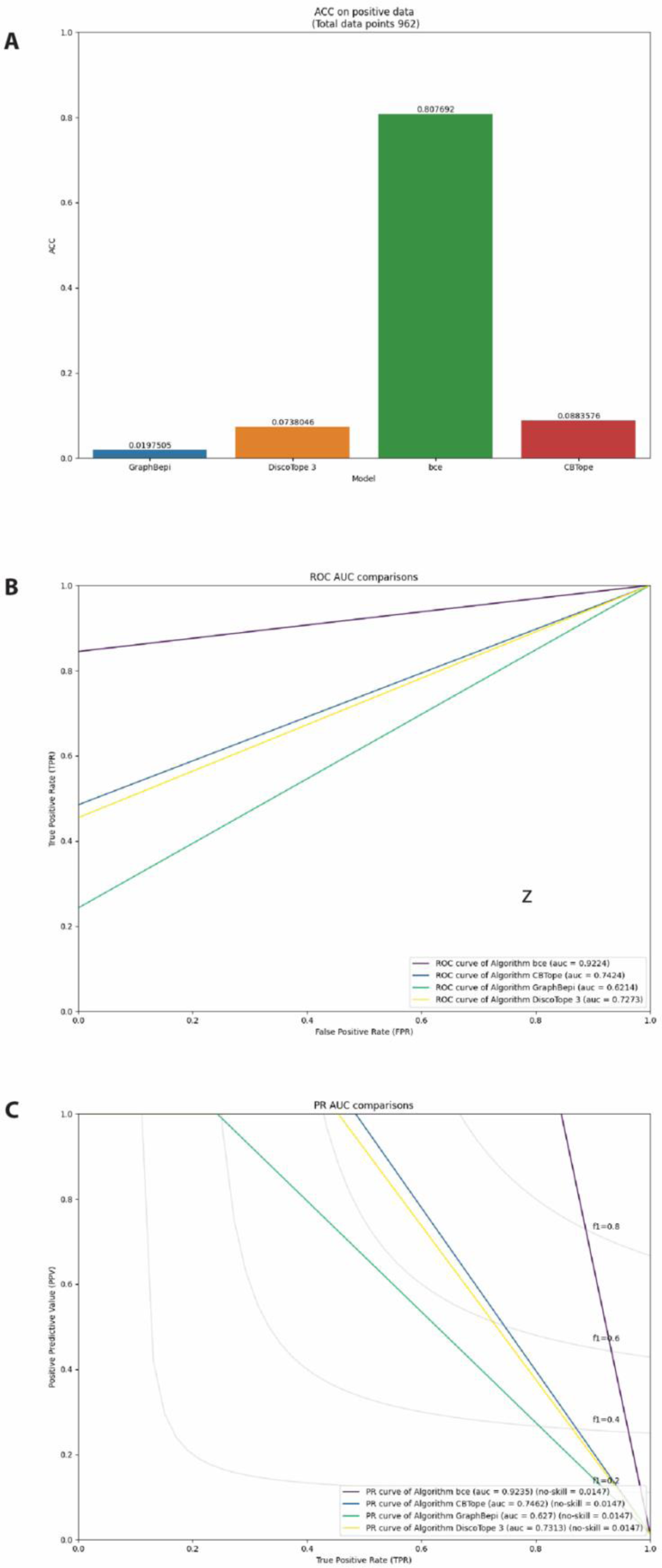
Benchmarking against existing state-of-the-art BCE predictors. 3A highlights the superior performance of BCE-HUNT in comparison to existing conformational BCE predictors, using a criterion for success conceptually similar as to how an epitope is defined as MPVs in BCE-HUNT. 3B and 3C highlights the significantly improved performance of BCE-HUNT compared to existing tools when using the criterion for success as a measure of the existing tools capability to predict at least on AA in the epitopes, for ROC and PR AUCs respectively. In both types of analyses (in 3A and 3B/C) BCE-HUNT must, strictly, predict the entire epitope (each AA in the epitope must be predicted correctly for directly physically interacting with the antibody and not directly interacting).

Since we only have true positives (TP) in the independent test of experimentally verified 3D antibody-bound protein structures, we used accuracy as the percentage of correct classifications (ACC) that each model in the comparison makes, as the evaluation metric 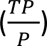. An explanation for how this evaluation metric is derived is outlined in the methods section. The outcome of this first type of evaluation is summarized in Figure 3A, where it is clearly demonstrated that BCE-HUNT significantly outperforms the existing tools, based on an evaluation that is conceptually similar as to how an epitope is defined as MPVs.

In the second type of evaluation against the existing state-of-the-art tools, we devised an experimental comparison that was conceptually similar as to how an epitope is defined on a per AA basis, like these existing tools. That is, if one of the existing tools identifies an AA on the experimentally verified epitope, then that is counted as a TP positive success. For example, considering again the >0.9 threshold for Discotope3.0 (32), if a true epitope in the test is [**1**,0,0,**1**,0,**1**] and if, for example, Discotope3.0 predicted that an AA contributes to the epitope if it has a score in the output of its model of > 0.9, the following was counted as a successful TP for Discotope3.0 if it has the following score:[0.6,0.02,0.3,**0.98**,0.2,0.7] (even though it has predicted only one of the AA acids positively in the entire epitope). For BCE-HUNT, in this analysis we enforced the strict criteria whereby 100% of the epitope experimentally verified to be physically participating in the interaction must be predicted correctly. To explain this in other terms, if there are 3 AAs contributing to the epitope in the MPV, BCE-HUNT captures is considered to predict either 0/3 AAs (failure) or 3/3 AAs (success) for any given epitope being tested in the AUC calculations. Whereas the other tools were allowed to score parts of the epitope (1/3, 2/3 or 3/3 AAs) and counted as successful positive hits in the AUC calculations. When BCE-HUNT did not predict 100% of the epitope correctly (3/3), the prediction was strictly counted as a failure for BCE-HUNT in the computation of ROC-AUC and PR-AUC curves in Figure 3B and Figure 3C, respectively. It was interesting to observe in Figures 3B and 3C, that although the relaxed criterion (applied to the second type evaluation) was more favorable to the per AA epitope predictors, BCE-HUNT outperforms the existing tools based on both PR and ROC AUC metrics.

For all the tools tested the threshold score for an AA was taken to be the default score as described in the respective published study for that algorithm (31–33). The threshold for a true positive hit for an AA in an epitope for BCE-HUNT was very strict in these comparisons (for both types of evaluation); a probability of being an epitope score greater than 0.96 was required to be deemed positive 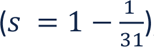, given the 30X ratio of positive to negatives in the training data in BCE-HUNT.

### Validation based on experimentally verified SARS-CoV-2 spike protein antibody epitopes and dissimilar experimentally verified antibody-bound protein 3D structures

We next proceeded to assess the performance of the BCEP-HUNT model under a useful case example whereby artificial intelligence (AI) guided B cell antibody epitope mapping could potentially hold promising biomedical benefits. Specifically, we assess the ability of the model to identify *bona fide* epitopes on the spike protein of the SARS-CoV-2 virus. This was a highly relevant validation-case example due to the recent surge of studies that have characterized the neutralizing antibody interactions with the receptor binding domain of the spike protein since the declaration of the COVID-19 pandemic in March 2020. We extracted 177 spike protein-antibody complexes from experimental sources (such as X-ray crystallography or cryo-electron microscopy from the PDB, see Supplementary Table 1 for the list of PDB Ids tested). None of the PDB complex structures used in this test were present in the training data for BCE-HUNT. In turn we then extracted 312 epitopes from these complexes. In Figure 4 we demonstrate the ability of BCE-HUNT to correctly predict the majority of these 312 epitopes. At a very strict threshold score of 0.9 for BCE-HUNT we were able to successfully predict 64% of these epitopes, and 73% at the more relaxed score of 0.5 (Figure 4A). To gauge how the state-of-the-art existing tools would perform on the same data, we chose the best performing tool from the independent benchmarking in Figure 3, CBTope (33), demonstrated that the best performing state-of-the-art tool could only predict a mere 0.01% of the BCEs (see Figure 4B). In each epitope test, a strict criterion was forced on BCE-HUNT in that it had to predict the entire epitope as a perfect match.

**Figure 4:**
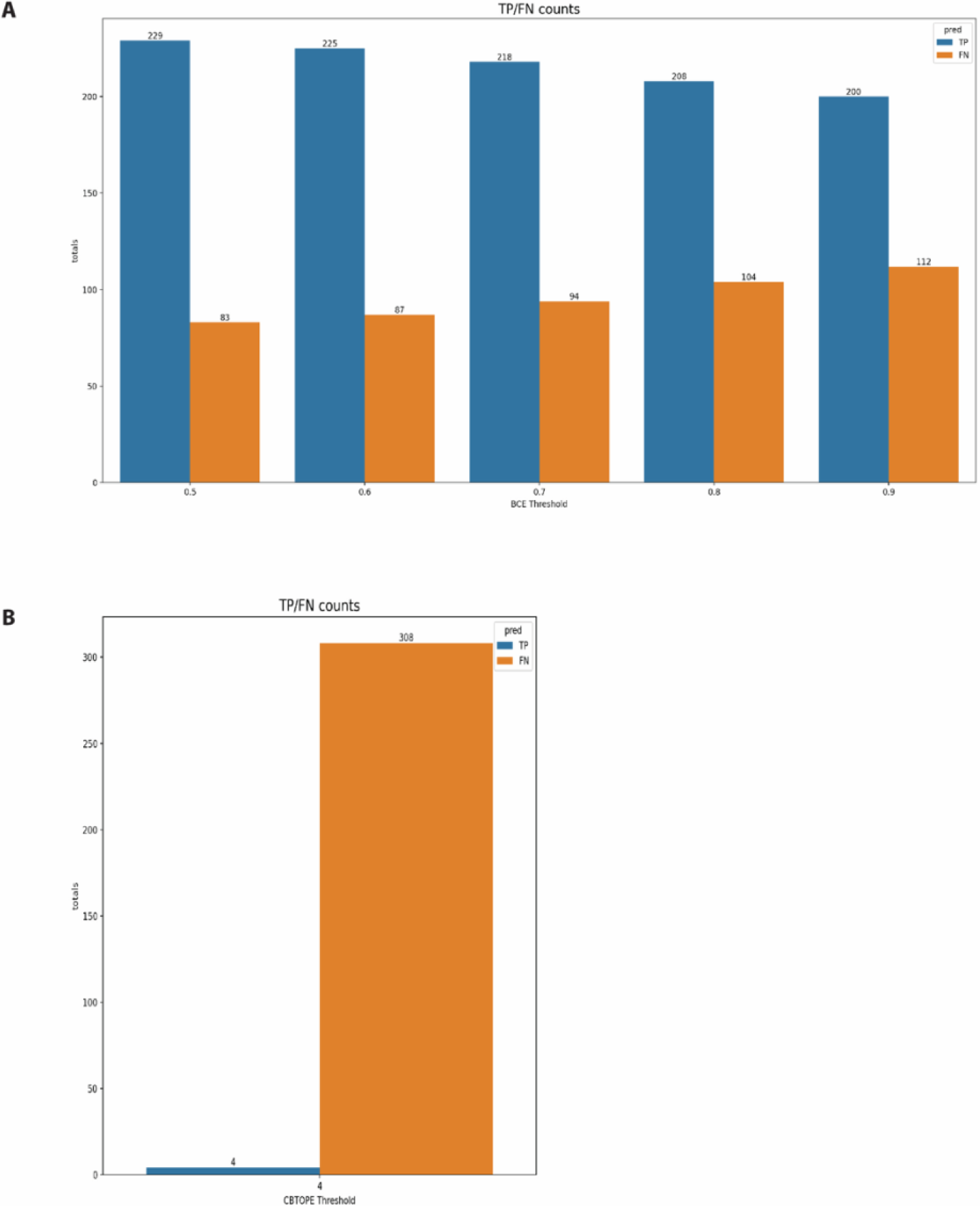
Validation on unseen SARS-CoV-2 spike protein antibody bound complex structures. (A) BCE-HUNT was successfully capable of recovering between 64% to 75% of bona fide epitopes experimentally validated antibody bound spike protein complexes (ranging from BCE-HUNT positive hit thresholds from 0.5 to 0.9). None of the antibody-spike protein complex in this test were included in the training data for BCE-HUNT, and the model needed to predict the epitope with a perfect match in the structural properties to be counted as a positive hit. (B) Using the same antibody bound SARS-CoV-2 spike test from the best performing state-of-the-art tools in Figure 3, CBtope, it was clearly demonstrated that the current approaches to predict BCEs are not fit for purpose compared to the strategy outlined here.

The superior performance of BCE-HUNT to capture experimentally validated unseen BCEs on the spike protein of SARS-CoV-2 does potentially give rise to the supposition that BCE-HUNT performed so well due to antibody-bound spike protein structures from coronaviruses present in the BCE-HUNT training data (albeit, also likely for the existing state-of-the-art tools). Therefore, we next assessed if BCE-HUNT can successfully predict experimentally validated BCEs that are not only unseen previously by the model as shown in Figure 4, but also dissimilar to the protein sequences that exist in the training or test data. To perform this strict test, we fetched all data from the PDB that was not present in any of our previously used training or test data sets. Global alignments on these sequences were performed using Needle-EMBOSS (34) to identify dissimilar proteins at various cutoffs of sequence coverage identity. A protein sequence was kept if its sequence identity was equivalent to or lower than at least one of the protein sequences in all the training data for the given threshold % identity coverage (see Supplementary Table 2 for the final list of antibody-bound 3D protein structures used for this test). Figure 5 illustrates that BCE-HUNT can generalize and successfully predict on unseen data in a robust manner. For example, at the strictest threshold for the probability of being a positive BCE (>0.9), and at the strictest threshold of identity coverage (20%); BCE-HUNT successfully predicted 51% of the true positive epitopes (Figure 5). The best performing existing state-of-the-art tool, CBTope, predicted only 2% of the true positive epitopes at 20% threshold for sequence identity.

**Figure 5:**
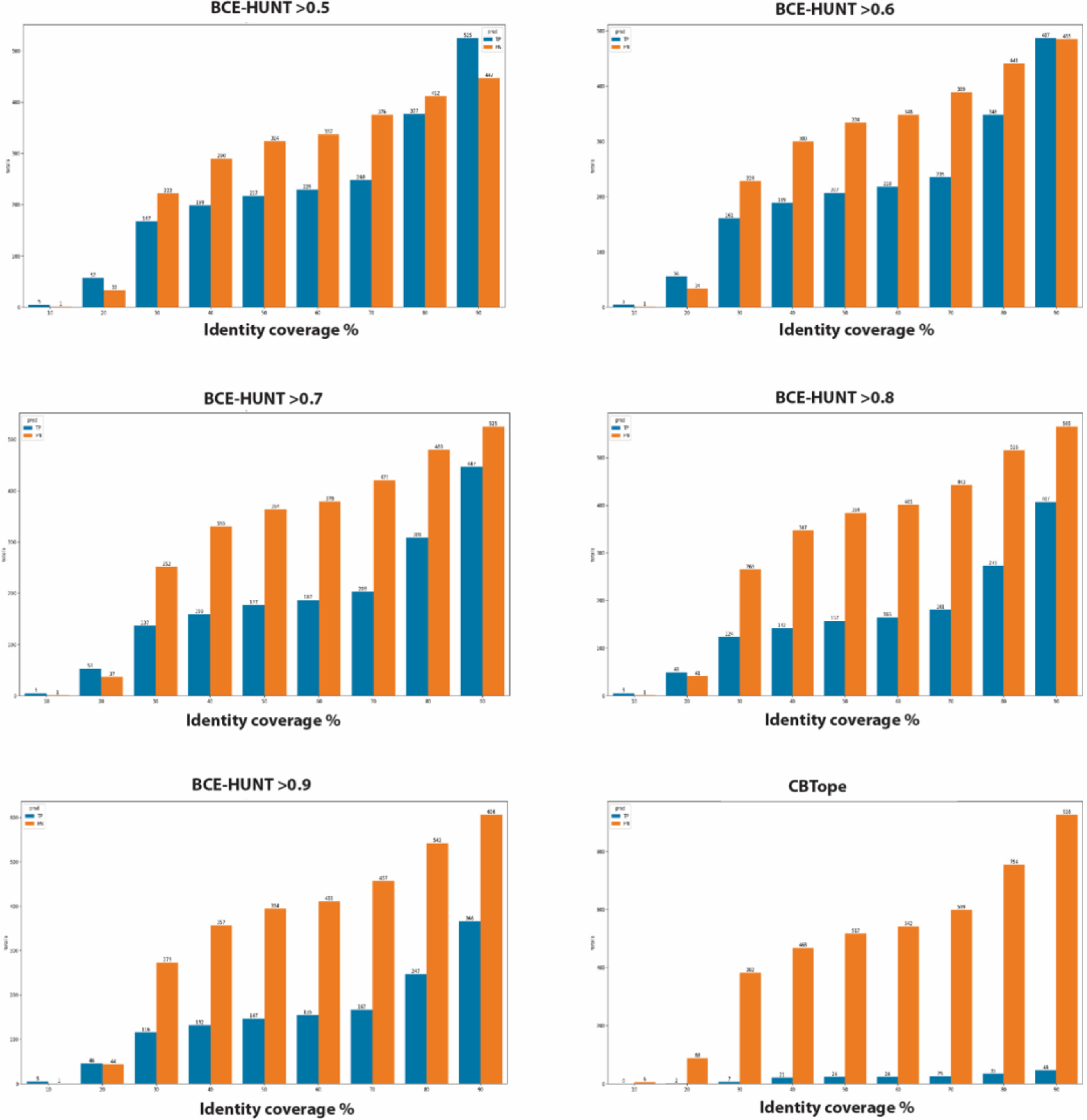
Validating against unseen or highly dissimilar BCEs from antibody bound protein 3D structures. Each plot shows the true positive (TP) epitopes that are identified as such from the predictor, on the given probability threshold (BCE-HUNT positives, ranging from positive probabilities of 0.5 to 0.9), and the false negative (FN) epitopes that are not identified as such from the predictor, on the given probability threshold. The X-axis depicts the identify coverage % threshold cutoff of the test data. Each threshold on the X-axis represents the population of BCEs on experimentally verified antibody-bound protein structures that are below the given threshold, and both unseen by the model and dissimilar to all the protein sequences in the training data. This analysis demonstrates the ability of BCE-HUNT to generalize and successfully predict on unseen data in a robust manner. This performance was far superior to the best state-of-the-art tool analyzed in Figure 3, CBTope.

## DISCUSSION AND CONCLUSIONS

A high performing machine learning (ML) model that accurately predicts BCEs can identify vaccine diagnostic candidates for further experimental validation much faster and more efficiently than experimental approaches alone. Such models can vastly reduce the cost and time related to the BCE mapping process and streamline the identification of potential epitope candidates for further clinical investigations.

Many approaches to predict BCEs have been attempted (7, 10, 35–39), but their performance is poor and there is significant room from improvement. Despite the development of several BCE predictors in recent years, computational BCE predictions have a long way to progress toward their practical use to guide the development of vaccines, antibody-based therapeutics and diagnostics. This was notable during the recent COVID-19 pandemic, whereby a wealth of additional ML training data became available due to the large burst of experimental studies on the spike protein from the SARS-COV-2 (38). This increased volume of spike glycoprotein structures, including antibody-bound structures, together with the more than 20 years of development of BCE predictors, could have potentially contributed significantly to the AI accelerated design of vaccines or monoclonal antibody-based therapeutics, however, the field reverted to 3D cryo-electron microscopy to guide the design of antibody-based therapeutics (40).

This recent example suggests that there is ample scope for improvement of BCE predictors, and here we present a novel approach that encompasses several unique and distinguishing innovations that more accurately model the underlying biology of BCE/antibody recognition and improve performance. The first innovation is the use of a more structural definition of the BCE, which captures the entire protein antigen sequence in a binary vector, where 1’s represents direct AA contact points on the 3D antigen structure with the antibody, and 0’s represents AAs that are not interacting directly with the antibody structure. The second innovation is the use of the unbound 3D antigen structure (rather than the antibody-bound structure) as a source of features to capture the pertinent 3D structural properties of the antibody-BCE binding interaction (before the antibody binding event occurs) termed here as the 3D macrostructures.

A similar concept underlying the second innovation, using the unbound protein antigen 3D structure to capture features before the binding event, has been reported previously in the literature (41–46). However, these previous antibody bound/unbound approaches used features captured from the unbound 3D protein antigen structure to directly train their respective BCE predictor models, and their definition of BCEs was on a per amino acid basis (not using MPV type epitope definitions). In the approach proposed here, the unbound 3D protein antigen structures are used to train several distinct ML models that capture the 3D macrostructure of 3D folded proteins (*e.g.,* RSA, HSE, and SS). Using the target antigen protein sequence alone, the outputs from the trained 3D macrostructure models are in turn used as features to train our main BCE predictor model (BCE-HUNT). Coupled together, both innovations outlined in the proposed approach, when combined, offer significant performance improvements compared to current state-of-the-art approaches. One of the limitations of the proposed approach is the fact that the user must query an almost unlimited number of MPVs to fully map the complete BCE potential in any sufficiently large antigen structure. This problem is currently handled by using a brute force approach where a random selection of hundreds of thousands of candidate MPVs, guided by priors from empirical BCE data are used, which are hopefully relatively representative of the broader BCE potential. However, this approach will undoubtedly miss good candidates and in future versions of this technology, we propose the integration of deep reinforcement learning (DRL) strategies to identify an even more representative and optimized space of BCEs represented by MPVs among the innumerable possibilities in any given protein antigen. In recent years, BLSTMs have been used to capture the global properties that may define the landscape of BCE in a protein antigen (31, 47). The use of BLSTMs have a key advantage as they allowed us to capture global relationships between distant amino acids which might constitute a conformational BCE. In contrast, other algorithms used in the other tools described earlier segment the protein sequences and consequently risk losing these relationships. However, the previous BLSTM approaches, and most other approaches limit the identification of BCEs on a protein to outputting a score on a per amino acid basis. The proposed advantage of the BCE as a binary MPV is that the structural properties of the epitope-paratope interactions can be captured with improved fidelity. Even though, BCE-HUNT can also be adjusted to perform predictions on a per amino acid basis (as performed in this study for benchmarking purposes), the MPV definition of a BCE is preferential as it encodes a high-resolution definition of each amino acid contribution in the epitope leading to significantly higher performance as reported here.

Another, advantage of the approach we describe herein is that it circumvents the need for an experimentally validated 3D structure to be available at the input stage, which is required by other approaches described earlier such as the DiscoTope family of BCE prediction tools (32, 48, 49). A recent development (32) in the BCE prediction field to address the limitation of requiring the PDB 3D protein structure as input, is to integrate the input query with the input 3D protein structure from databases such as the Alphafold protein structure database (50). Although the Alphafold method (51) is indeed a ground-breaking advancement in 3D protein structure prediction, and serves as an important hypothesis generator, 3D protein structure predictions are arguably not yet a direct placement for experimental structure determination (12). Therefore, strategies such as BCE-HUNT described here are advantageous in that they require the protein sequence alone to perform reliable BCE predictions. As mentioned above, BCE-HUNT bypasses they need for the experimentally derived 3D protein structure or potentially unreliable predicted 3D protein structure at prediction-time, by capturing the key 3D macrostructure properties of the protein from the unbound protein antigen during ML training in the pipeline. We demonstrated in this study that the precise coordinates of each atom in each amino acid relative to each other the 3D protein structure is not necessary to be known with fidelity to capture accurate BCE antibody interactions. The determinants of the 3D protein structure captured by the 3D macrostructure predictors from the unbound protein antigen may be sufficient.

Given the failure of the existing tools to reliably predict BCEs on the spike protein of SARS-Cov-2 (7), the validation case study we report herein on unseen SARS-Cov-2 antibody bound spike protein structures suggests that the approach proposed in this study offers an important advancement in the field and may help pave the way towards a future where computational BCE prediction is routinely used in a wide range biomedical research applications and to help design future vaccines and immunodiagnostics.

## METHODS

### Pretraining techniques

For ensuring reproducible results and avoiding the need of random seed in the networks used in our BLSTM models, we used auto-encoders (AEs). AEs are types of neural networks which mirror their inputs. More specifically, an AE takes an input at layer number i = 0 and processes it through an arbitrary number of layers, say i = 1, …, N, which constitute the encoder part. It then processes it back through a mirrored structure of the encoder part, called the decoder part. Finally, it returns the output which is the same as the input. The intermediate layers might reduce or expand the dimensions of the input, as the main use of AEs was indeed dimensionality reduction. Firstly, we randomly generated 10000 protein sequences of length between 50 to 1000 amino acids. The actual amino acids used was random as well. We then pre-trained each layer of the neural network as an individual AE on the whole generated data. The pre-training was done for maximum 1000 epochs for each AE, with the possibility of early stopping if the loss did not decrease after 10 epochs. The weights of the outer layers were copied to deeper AEs after pre-training and were also kept constant during the pre-training of those. The last layer of the model (output) was not pre-trained at all. This procedure was done once, and the same pre-trained weights were used on all the validation procedures. This procedure was faster, it used much more data to pre-train and the same initial weights were used for all downstream analyses.

### BLSTMs

Bidirectional long short-term memory networks (BLSTMs) consist of two LSTMs, each one scanning the time-steps sequence from either direction. That is, one LSTM scans the forward and one the backward sequence of time-steps. This allows the network to capture relationships between past and future time-steps at once. For predicting conformational BCEs, we use BLSTMs models. This allowed us to model the whole protein sequences as one observation, without the need of segmenting it. Therefore, distant amino-acid relationships should be able to get captured by the model. Moreover, the use of BLSTMs allows us to train different protein lengths simultaneously.

### Data preparation for the 3D macrostructure features (unbound 3D protein structure data preparation)

To accurately predict the 3D macrostructure features (SS, RSA, UHSE and LHSE) from a native protein sequence, we used 3D protein structures from the PDB(13) from all organisms. The goal of those models was to predict the surface and structural characteristics of proteins that are not affected by any other protein, including antibodies (Abs). Therefore, we kept only structures that are not bound by any other structure or molecule. Structures containing more than one copy of the same molecule, but slightly different conformation, are kept in the data. We filtered the structures and kept only those with ≤ 3°A resolution, ensuring that every atom of each amino-acid is mapped with coordinates, and with protein chains longer than 200 amino-acids. After filtering, the database consisted of 41592 total structures (per 20/12/2019). Those structures contained 70489 protein sequences which resulted in 53524 unique sequences. Subsequences of longer sequences were kept as different data points. The reason for that is that their conformational characteristics might be different because of their shorter length.

The DSSP algorithm was used to compute the SS and the RSA for each molecule in each structure file (52). DSSP computes the following secondary structure classes for a protein sequence; α-helix, 310-helix, π-helix, isolated β-bridge, β-strand, turn, bend and coil. We merged those classes into three super-classes; Helices (α-helix, 310-helix and π-helix), Strands (isolated β-bridge and β-strand) and Coils (turn, bend and coil). Finally, the BioPython package (53) was used to compute the UHSE and LHSE. At the end of the filtering, each amino acid in the data base was assigned a value for the RSA, UHSE, LHSE and a class for the SS. To create a unique data base, the mean per amino-acid was taken for RSA, UHSE, LHSE among identical sequences. Finally, amino-acids of identical sequences but with different SS classes, were assigned to the coil class.

### BLSTM Prediction model for RSA and HSE (3D macrostructure features)

We chose to create a single BLSTM model for predicting all surface features. More specifically, the model predicts RSA, UHSE and LHSE from a primary protein sequence. Although LHSE might not give useful information about the surface position of an amino acid, it might help predicting UHSE more accurately, since both together are actually forming a prob around each amino-acid. This is a BLSTM model which takes as inputs a batch of features, each computed per amino-acid from each input sequence, and predicts a three-way output. For each protein sequence given as input, a value for each RSA, UHSE and LHSE are predicted per amino-acid. For all the three outputs the individual losses were the mean square error (MSE). The global model loss was the weighted sum of the individual losses with weights: 50, 100 and 125 for RSA, UHSE and LHSE, respectively. The weights were decided by first training the model without them and see the differences in the magnitude of the three losses. The weights contribute in such a way that the three losses give the same contribution to the global model loss. Supplementary Figure 2A shows the network architecture of the model, and Table 1 outlines the features used to the train the BLSTM model.

### BLSTM Prediction model for secondary structure (3D macrostructure features)

We also chose to create a BLSTM model for predicting a three class output for SS.The model uses the categorical cross-entropy loss in order to assign one of the following classes to each amino-acid in an input sequence; Helices, Strands and Coils. Supplementary Figure 2B shows the network architecture of the model, and Table 1 outlines the features used to the train the BLSTM model.

### Data preparation for the conformational BCE predictor (BCE-HUNT)

For modelling conformational BCEs (CBCE) we downloaded non-obsolete protein complexes from the PDB(13). We allowed structures of any resolution and organisms, if they had at least three different protein chains. The reason for this is that two of the chains might be the variable fragment heavy (VH) and variable fragment light (VL) chains of an antibody (Ab), and the third chain might be an antigen (Ag). Of course, there might be multiple Abs or Ags in one PDB structure. We created a local database using all immunoglobulin V-, D- and J-region genes from the international ImMunoGeneTics information system (IMGT)(54). Those genes were downloaded for both VH and VL chains, from all the available organisms, that is, human, mouse, rhesus monkey, rabbit, and rat. The IMGT genes not only provide information about the VH and VL chains of an Ab, but its paratope regions as well, that is, the complementarity-determining regions (CDR) and framework regions (FR) of each chain. To identify the Abs, we used IgBlast (55) for protein sequences on the IMGT database. For identifying the paratope in a chain we used the Kabat system that the IgBlast provides (56). We considered VH and VL chains as valid, only if at least their CDR1 and CDR2 were found by IgBlast. Since CDR3 is more difficult to map (57), we allowed it to be missing. We blasted all the protein chains of each structure to that database. Structures were filtered out if they did not have at least one chain mapped as valid VH and one as valid VL. Any other chain that was not mapped as VH or VL was assumed to be an Ag chain.

We considered as valid Ag chains those chains that were longer than 100 amino acids and were not bound by any other chain other than a VH or VL chain. Finally, only structures including at least one VH, VL and Ag chains were taken to further analyses. We paired VH and VL chains in each structure to recreate valid Abs. In case there were multiple VH and VL chains in one single structure, we measured the mean distance from every atom on each VH to every atom on each VL chain. VH and VL chains with the minimum mean atom distance were assigned as pairs and assumed to belong to one single Ab. Stand-alone VH or VL chains were not considered on the downstream analysis as they could not define a complete Ab. A paratope analysis provided information about the contacts formed with each Ag. Two amino acids were assumed to be in contact if any of their atoms were located within a certain probe distance from each other. We computed the total number of Ag amino acids that form contacts with each paratope’s parts, using probe distances of 4, 6 and 8 °A. Most of the contacts are made with CDR1 and CDR2 of the VHs and CDR1 and CDR3 of the VLs. The absence of CDR3 on VHs might be due to the difficulty of identifying it using IgBlast. Moreover, the FR regions do not seem to form as many contacts as the CDR regions, as expected. Additionally, the standard deviation of the total contacts per amino acid and paratope part was relatively low, indicating that similar number of contacts are made between different Ags and Abs. This could be an indication of the possibility of predicting CBCEs without any further information about the actual Abs. For each VH and VL chain pair we defined a single CBCE. This was done by first identifying atoms on any Ag whose distance from the CDRs regions of any VH and VL chain pair was ≤ 4°A. The amino acids that those atoms belonged to were defined as contacts between the Ag and the Ab, that is, they defined the CBCE. Multiple CBCEs from different Abs could be defined on the same Ag. Finally, structures that did not define any CBCE within the 4°A distance were discarded. Observed CBCEs were also mapped to similar Ag. Undiscovered CBCEs could increase the false negative rate of the prediction models. To decrease the possibility of assigning undiscovered CBCEs as negative data, we copied observed CBCEs to similar Ags. First, we identified clusters of similar Ags using BlastPlus (58) with > 90% similarity and at most two gaps. CBCEs were copied from an Ag in a cluster to all the other Ags in the same cluster if their corresponding position and distancing was the same based on the mapping from BlastPlus. Lastly, we created a unique Ags database. Duplicated Ags were removed from the data. Second, Ag sequences that were sub-sequences of longer Ags were kept in the data and treated as different observations. The resulting database consisted of 1003 Ag sequences in fasta format and 12968 CBCEs from which 6986 were unique. Each Ag sequence was associated with at least one CBCE.

### BLSTM Prediction model for the conformational BCE predictor (BCE-HUNT)

The models utilize input features that include, among others, a permutation vector, also known as a molecular permutation vector (MPV). Every sequence in the data is associated with at least one true CBCE, those CBCEs are turned into binary 2D vectors and are given as input to the models. The model’s primary output is a probability, which essentially addresses the query: “Does this particular permutation vector within the given specific sequence accurately constitute a true CBCE?”. The second output of the model is the permutation vector itself. This part of the model works as an AE, where it returns a probability per amino acid. This output can be seen as a contribution of each amino acid in the sequence to the specific CBCE in question. The goal was to predict CBCEs before the binding event took place. The dataset used comprises linear protein sequences and the true CBCEs. Therefore, it lacks information concerning the structural or surface characteristics of these protein sequences. However, an understanding of the pre-binding secondary structure and surface of each protein is crucial for our analysis. Therefore, we used our prediction models for RSA, UHSE, LHSE and SS (3D macrostructure features) to predict those characteristics in every protein in the dataset. Supplementary Figure 2C shows the network architecture of the BCE-HUNT model, and Table 1 outlines the features used to the train the BLSTM model.

The BCE AA output is the actual permutation output, where binary cross-entropy loss was used. The BCE perm output is a probability vector indicating if the input permutation is a true CBCE (second position on vector) or not (first position on vector). The input features were computed per amino acid as they are, not averaged by windows. The BCE perm output is computed from amino acid values. The last layer of the output is a Dense layer with sigmoid activation. Such that, for each protein sequence, a value in ∈ [0, 1] is returned per amino-acid. Then a 2-class probability vector is computed for that sequence as

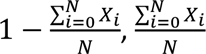

where N is the length of the sequence and *X_i_* is the dense layer output on amino-acid at position i on the sequence. This 2-class probability vector is then used in the binary cross-entropy loss.

### Negative data generation for the conformational BCE predictor (BCE-HUNT)

We applied three different negative data generating methods. For each sequence in the training and validation data we generated 30 completely random permutations, 10 from each method, and assumed that they are not true CBCEs. In the first method we generated completely random permutations. For each sequence in the training and validation data we generated 10 completely random permutations and assumed that they were not true CBCEs. Both the total number of amin-acids and the placement of those in the given sequence were random. New permutations were generated on each epoch. The advantage of this method is that because of the complete randomness, it is quite unlikely that any generated CBCE permutation will be false negative. However, the disadvantage is that the randomly generated CBCE permutations might be extremely different than the true CBCE permutations. In that case, the algorithm might learn to separate only based on the permutation input. The second and third methods correct for this disadvantage, which we also applied on every true CBCE of the given protein sequence and generated on each epoch. The second method kept the first and last amino acid of a given true CBCE at the correct position, while it randomly shuffled the internal CBCE amino acids indicators inside the region. The third method kept the total amount of amino acids of a true CBCE, as well their linear distance, constant. It then randomly shifted the whole true CBCE on other parts of the protein. The advantage of this method is that the randomly generated CBCE permutations cover both extremely similar and extremely different true CBCE permutations. This is likely to result in an algorithm that is more robust to both positive and negative data. Conversely, a drawback of the method is the potential for a substantial rise in false negatives, which could lead to diminished performance when applied to novel data sets.

### Evaluation metrics

For the evaluations against the existing state of the art tools in Figure 3A, ACC was used as the metric, which was derived here as the true positive rate (TPR) 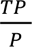. This was derived due to 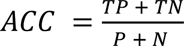. For those comparisons, we only used scientific verified antibody binding epitopes from the PDB. For that reason, and because we did not want to assume negative epitopes and risk FP rates, both the TN and N are zero. The ACC therefore in effect becomes the true positive rate (TPR), 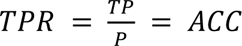.

For AUC calculations we assume that the 0’s representing AAs in the MPV are negative which allowed us to capture the TN rate in addition to the TP rate to perform the evaluations illustrated in Figures 3B and 3C.

