## Supplementary figures and images for "Deep learning of antibody epitopes using molecular permutation vectors"

### Supplementary Figure 1

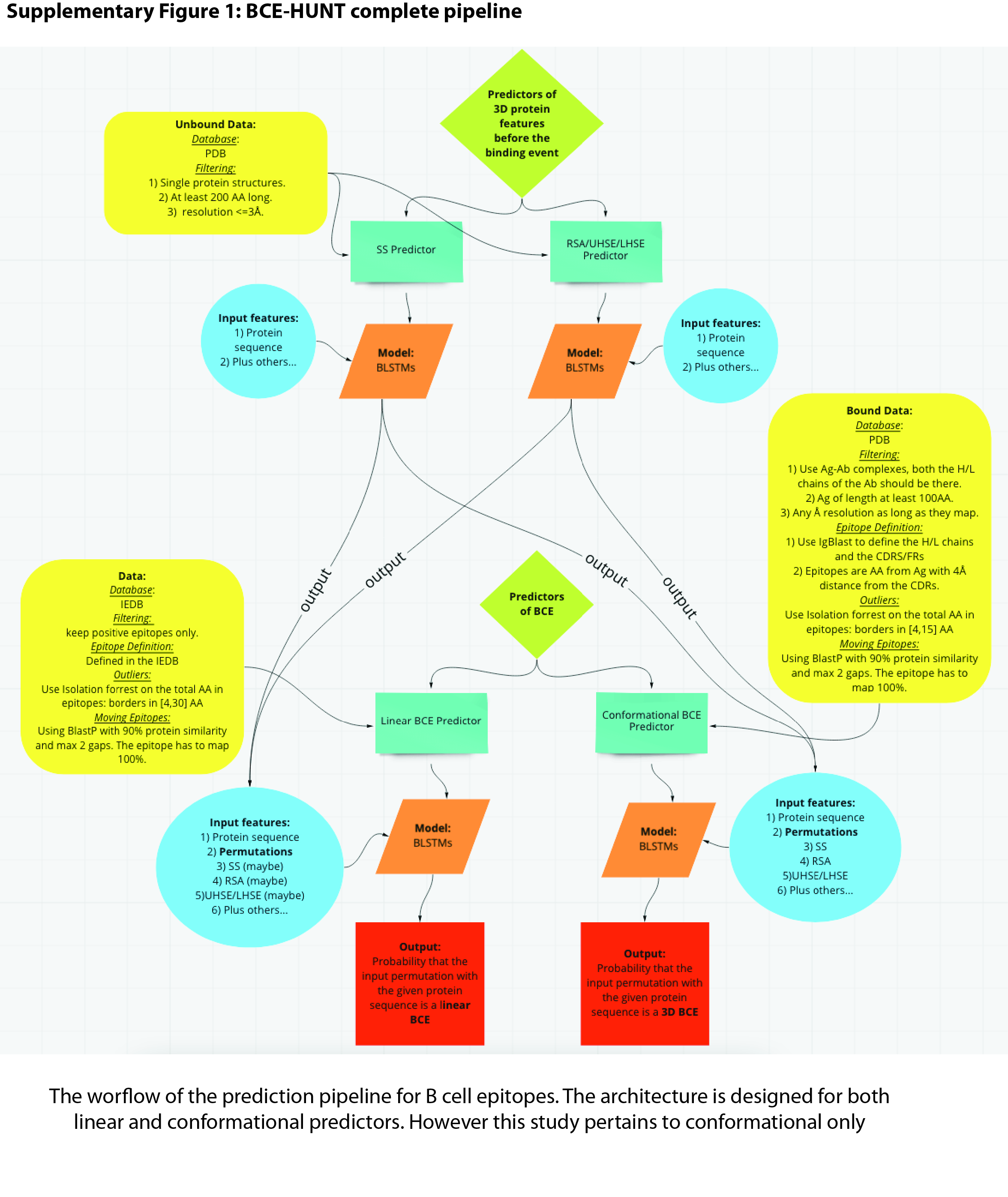
