## Supplementary Figure 2 for "Deep learning of antibody epitopes using molecular permutation vectors"

**Supplementary Figure 2: BLSTM Model architectures (3D Macrostructure models and BCE-HUNT)**

**A :** RSA, UHSE, LHSE BLSTM model architecture

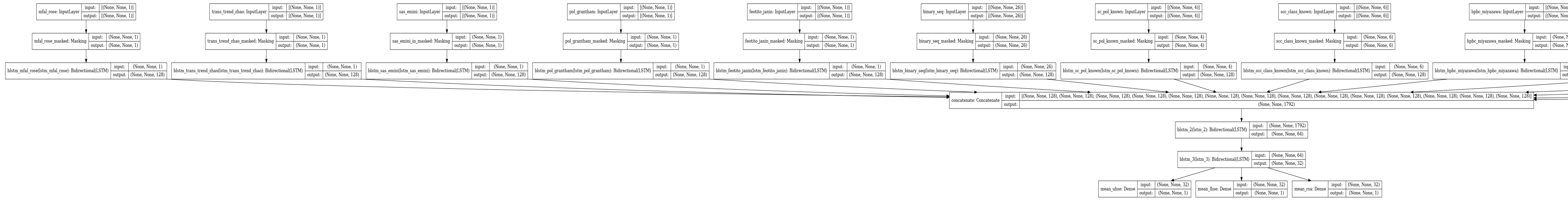

**B :** Secondary structure BLSTM model architecture

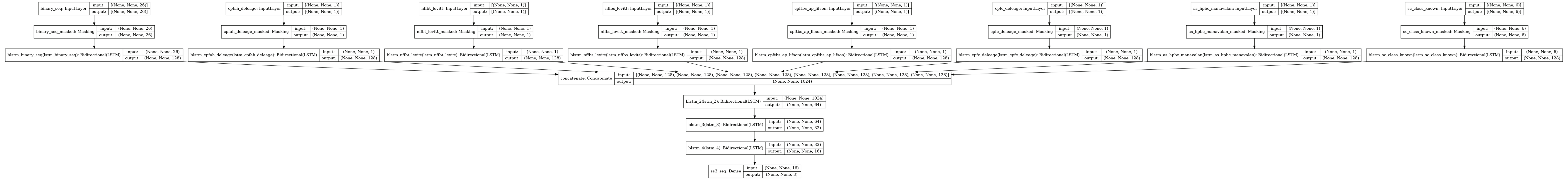

**C :** BCE-HUNT BLSTM model architecture (main BCEP predictor)

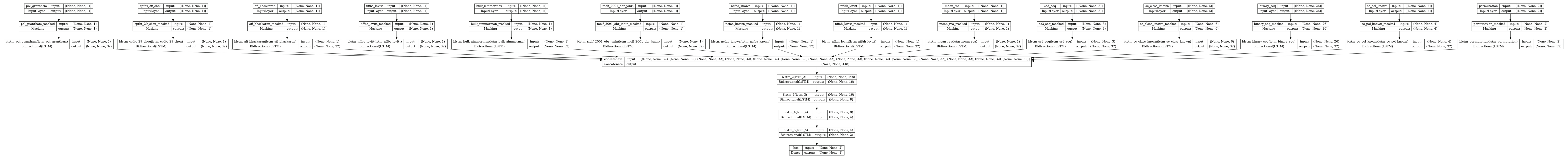
